# Genetically diverse uropathogenic *Escherichia coli* adopt a common transcriptional program in patients with urinary tract infections

**DOI:** 10.1101/595207

**Authors:** Anna Sintsova, Arwen Frick-Cheng, Sara Smith, Ali Pirani, Sargurunathan Subashchandrabose, Evan Snitkin, Harry L. T. Mobley

## Abstract

Uropathogenic *Escherichia coli* (UPEC) is the major causative agent of uncomplicated urinary tract infections (UTIs). A common virulence genotype of UPEC strains responsible for UTIs is yet to be defined, due to the large variation of virulence factors observed in UPEC strains. We hypothesized that studying UPEC functional responses in patients might reveal universal UPEC features that enable pathogenesis. Here we identify a transcriptional program shared by genetically diverse UPEC strains isolated from 14 patients during uncomplicated UTIs. Strikingly, this *in vivo* gene expression program is marked by upregulation of translational machinery, providing a mechanism for the rapid growth within the host. Our analysis indicates that switching to a more specialized catabolism and scavenging lifestyle in the host allows for the increased translational output. Our study identifies a common transcriptional program underlying UTIs and illuminates the molecular underpinnings that likely facilitate the fast growth rate of UPEC in infected patients.

## Introduction

Urinary tract infections (UTIs) are among the most common bacterial infections in humans, affecting 150 million people each year worldwide (1). A high incidence of recurrence and frequent progression to chronic condition exacerbates the negative impact of UTIs on patients’ quality of life and healthcare costs (2). Despite the magnitude of the problem, treatment remains limited by a strain’s susceptibility to available antibiotics, which are often ineffectual (3–5).

The major causative agent of uncomplicated UTIs is Uropathogenic *Escherichia coli* (UPEC), which is responsible for upwards of 70% of all cases (1). The majority of our insights into UPEC pathogenesis have been obtained through *in vitro* assays, cell culture systems, and animal models (6–9). While these studies have identified virulence and fitness factors that are important for UPEC infection, how these studies translate to human infection is not clear. As a result, we do not yet have a complete understanding of UPEC physiology in the human urinary tract. Moreover, the genetic heterogeneity of UPEC isolates, which carry diverse and functionally redundant virulence systems including iron acquisition, adherence, and toxins, further complicates our understanding of uropathogenesis (10–14). The different constellations of virulence factors and diverse genetic backgrounds raise the question of whether different UPEC strains vary in their strategies for pathogenesis.

Since defining conserved UPEC characteristics have proven elusive to comparative genomics strategies, we hypothesized that comparing functional responses in the context of the host may uncover disease-defining features. To that end, we directly examined UPEC gene expression directly from 14 patients with documented significant bacteriuria and presenting with uncomplicated UTI and compared it to the gene expression of identical strains cultured to mid-exponential stage in filter-sterilized pooled human urine. Despite the genetic diversity of the pathogen and the human hosts, we identified a remarkably conserved gene expression program that is specific to human infection, can be recapitulated in the mouse model of infection and bears all the hallmarks of extremely rapid growth rate. Based on extensive analysis, we propose a model where UPEC shut down all non-essential metabolic processes and commit all available resources to rapid growth during human UTI. Critically, our discovery of a common transcriptional program of UPEC in patients significantly expands our understanding of bacterial adaptation to the human host and provides a platform to design universal therapeutic strategies.

## Results

### Study design

To better understand UPEC functional responses to the human host, we isolated and sequenced RNA from the urine (stabilized immediately after collection) from fourteen otherwise healthy women diagnosed with UPEC-associated urinary tract infection. To identify infection-specific responses, we cultured the same fourteen UPEC isolates *in vitro* in filter-sterilized human urine (mid-exponential phase, 2-hour time point in Fig S1), and isolated and sequenced RNA from these cultures (study design and quality control is described in detail in Methods section). Phylogenetic analysis showed high degree of genetic diversity, as we identified strains belonging to 3 distinct phylogroups (Fig. S2). The majority of UPEC isolates (10 of 14) belonged to the B2 phylogroup, which is consistent with previously published studies (2, 13, 16–18). Although the majority (10 of 14) of patients had previous history of UTIs, we found no relationship between patients’ previous UTI history and bacterial genotype (Fig. S2). Moreover, the fourteen clinical isolates showed a wide array of antibiotic resistance phenotypes (Fig. S2).

### Virulence factor expression is not specific to infection

We first assessed the virulence genotype of the fourteen UPEC strains by looking at the presence or absence of a comprehensive list of known virulence factors, including adhesins, toxins, iron acquisition proteins, and flagella (9–14)(Fig. 1A). As previously reported (13), B1 strains appear to carry fewer virulence factors overall when compared to B2 strains, suggesting that UTIs can be established by UPEC strains with vastly diverse virulence genotypes. We then compared the levels of gene expression of these virulence factors following culture in filter-sterilized urine (Fig. 1B) to that during infection (Fig. 1C). Gene expression during infection and in urine showed strain-to-strain variability, which is consistent with previous reports (15). Virulence factors were expressed at similar levels in *in vitro* urine cultures and during infection. For example, iron acquisition systems were expressed regardless of experimental condition (Fig. 1B, C). Notably, virulence factor carriage varies greatly between UPEC strains and we did not discern any infection-specific gene expression among the virulence factors we examined (Fig S3).

**Fig. 1.**
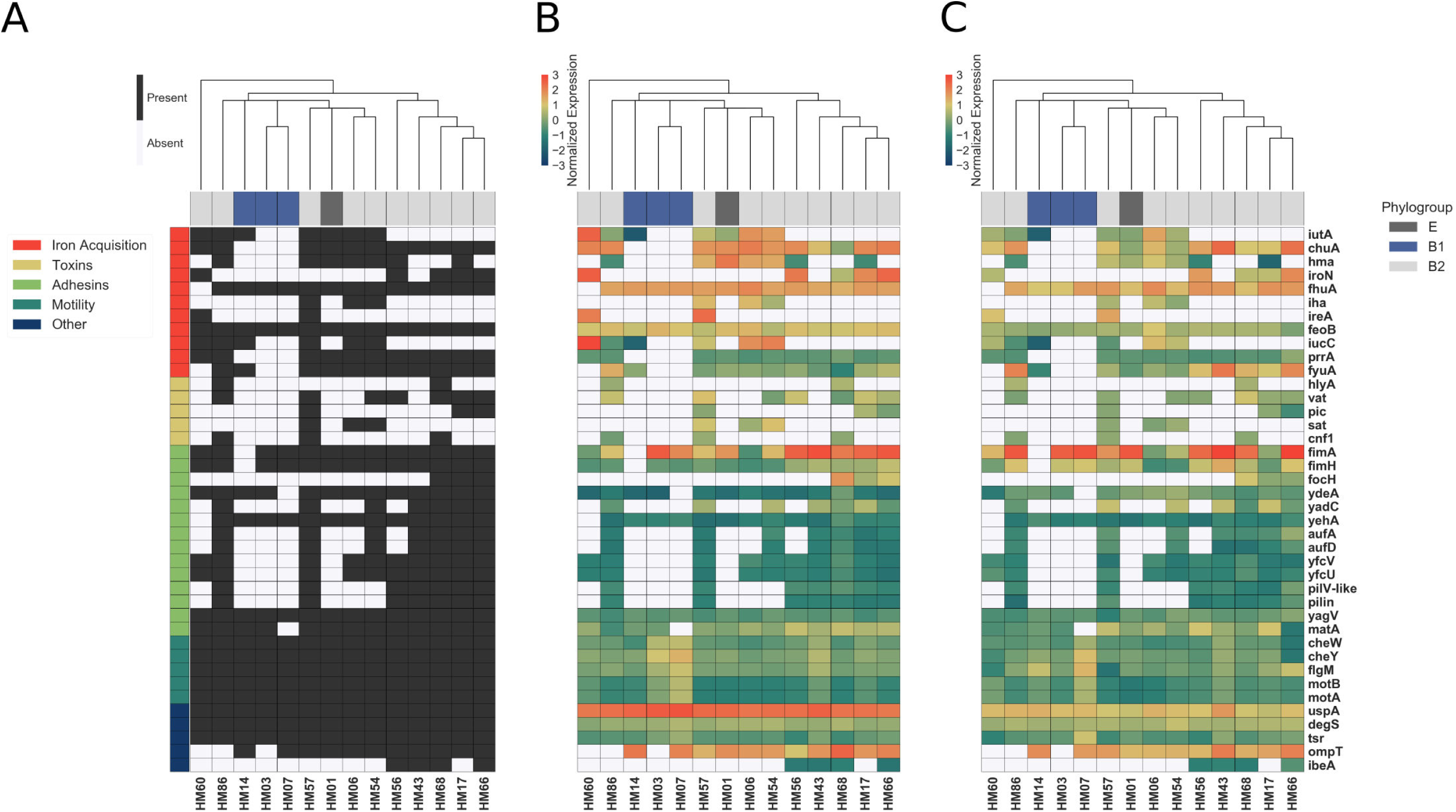
Clinical UPEC isolates carry a highly variable set of virulence factors. (A) Clinical UPEC isolates were examined for presence of 40 virulence factors. Virulence factors were identified based on homology using BLAST searches (≥80% identity, ≥90% coverage, average % identity indicated next to gene name). The heatmap shows presence (black) or absence (white) of virulence factors across 14 UPEC strains. Hierarchical clustering based on presence/absence of virulence factors shows separate clustering of B1 isolates. (B) Normalized gene expression of the 40 virulence factors in UPEC strains during *in vitro* urine culture. (C) Normalized gene expression of the 40 virulence factors in UPEC strains in patients.

### The UPEC core genome exhibits a common gene expression program during clinical infection

To perform a comprehensive comparison of gene expression between the different clinical UPEC strains, we identified a set of 2653 genes present in all 14 UPEC strains in this study as well as the reference *E. coli* MG1655 strain (hereafter referred to as the core genome; see Methods, Fig S4, Fig S5). We then assessed the uniformity of core genome expression of 14 isolates cultured *in vitro* in filter-sterilized urine. As expected for bacterial strains cultured under identical conditions, we saw high correlation of gene expression between any two isolates cultured *in vitro* (Fig. 2A, B (blue box), with median Pearson correlation coefficient of 0.92). Remarkably, the correlation of gene expression was just as high between any two patient samples (median of 0.91, Fig. 2A, B (red box)), while the correlation between *in vitro* urine and patient samples was considerably lower (median of 0.73 (Fig. 2A, B (green box)). The gene expression correlation between *in vitro* and patient samples remained low, even when we directly compared identical strains (*i.e.* HM56 cultured *in vitro* in urine vs. HM56 isolated from the patient, median of 0.74, Fig. 2A, B (yellow box)). This analysis suggested that UPEC adopt an infection-specific gene expression program that is distinct from UPEC undergoing exponential growth in urine *in vitro*. We confirmed this observation using principal component analysis (PCA), which revealed that patient samples form a tight cluster, distinct from *in vitro* cultures (Fig. 2C), demonstrating the common transcriptional state of UPEC during human UTI.

**Fig. 2.**
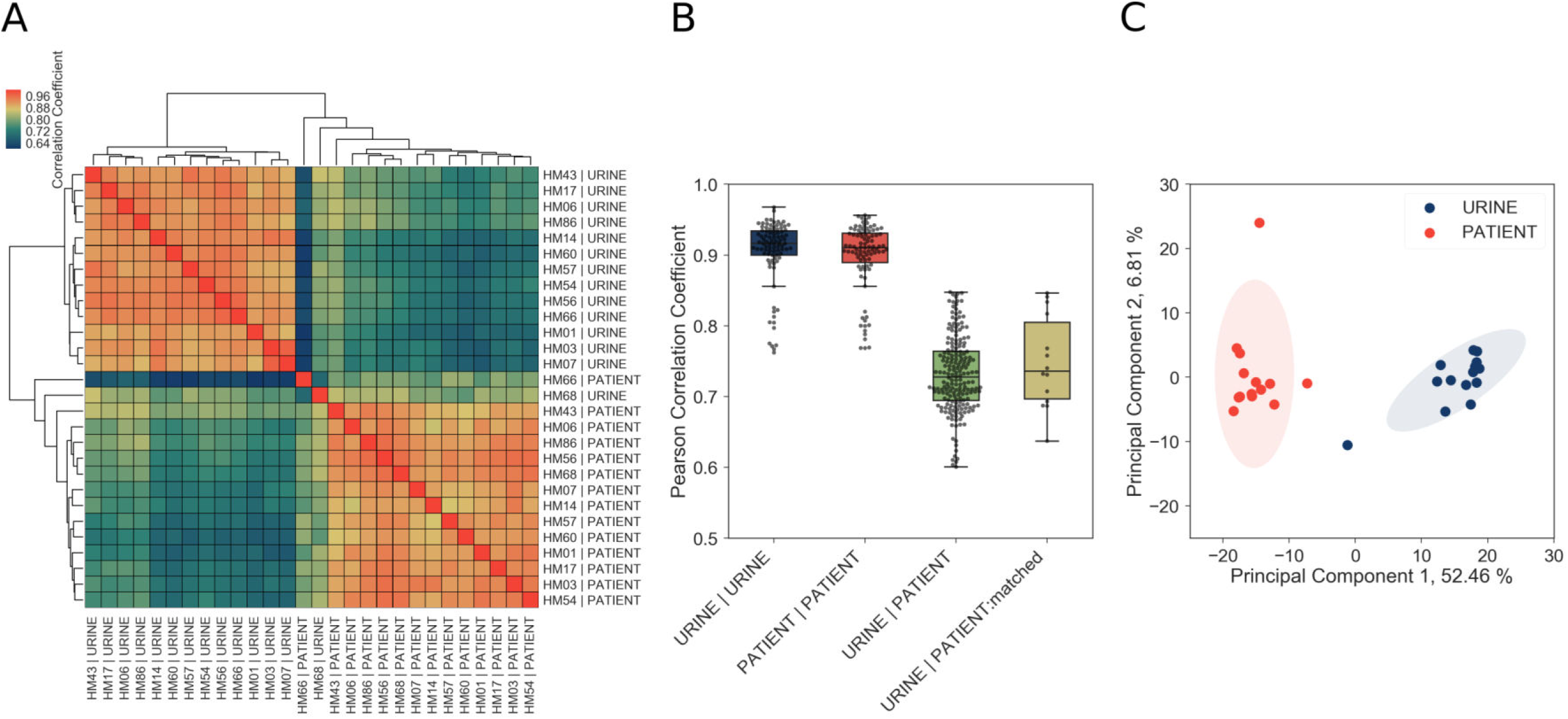
Core genome expression in patients is highly correlated. (A) Correlations among *in vitro* and patient samples measured by Pearson correlation coefficient of normalized gene expression plotted according to hierarchical clustering of samples. (B) Pearson correlation coefficient among all samples cultured *in vitro* (URINE | URINE, median = 0.92), among all samples isolated from patients (PATIENT | PATIENT, median = 0.91), between samples cultured in urine and samples isolated from patients (URINE | PATIENT, median = 0.73), and between matching urine/patient samples (ex. HM14 | URINE vs HM14 | PATIENT), (URINE | PATIENT:matched, median = 0.74). (C) Principal component analysis of normalized gene expression of 14 clinical isolates in patients and *in vitro* urine cultures shows distinct clustering of *in vitro* and patient isolates.

We also performed PCA analysis on *in vitro* (Fig. S6A, B) and patient samples (Fig. S6C, D) separately, to ascertain whether there is any discernable effect of bacterial phylogroup (Fig. S6A, C) or patients’ previous history of UTI (Fig. S6B, D) on gene expression. Interestingly, B1 and B2 strains did cluster separately and a number of genes were expressed differentially in B1 and B2 backgrounds (**Dataset S1, S2**), suggesting that variation in gene regulatory elements between phylogroups has a small but discernable role in gene expression both *in vitro* and during infection. However, we found that patients’ history of UTI had no effect on bacterial gene expression.

Taken together, our data indicate diverse UPEC strains adopt a specific and conserved transcriptional program for their core genome during human infection.

### UPEC show increased expression of replication and translation machinery during UTI

Differential expression analysis of the infection and *in vitro* transcriptomes identified 492 differentially expressed genes (log_2_ fold change greater than 2 or less than −2, adjusted *p* values < 0.05) (Fig. 3A, **Dataset S3, S4**). Interestingly, pathway analysis and manual curation of the differentially expressed gene list revealed that expression of ribosomal subunits (r-proteins), and enzymes involved in rRNA, tRNA modification, purine and pyrimidine metabolism, are significantly higher in patients compared to *in vitro* cultures (Fig. 3B, Table S2). Together, these data strongly suggest that replication rates during infection are significantly higher than during mid-exponential growth in urine.

**Fig. 3.**
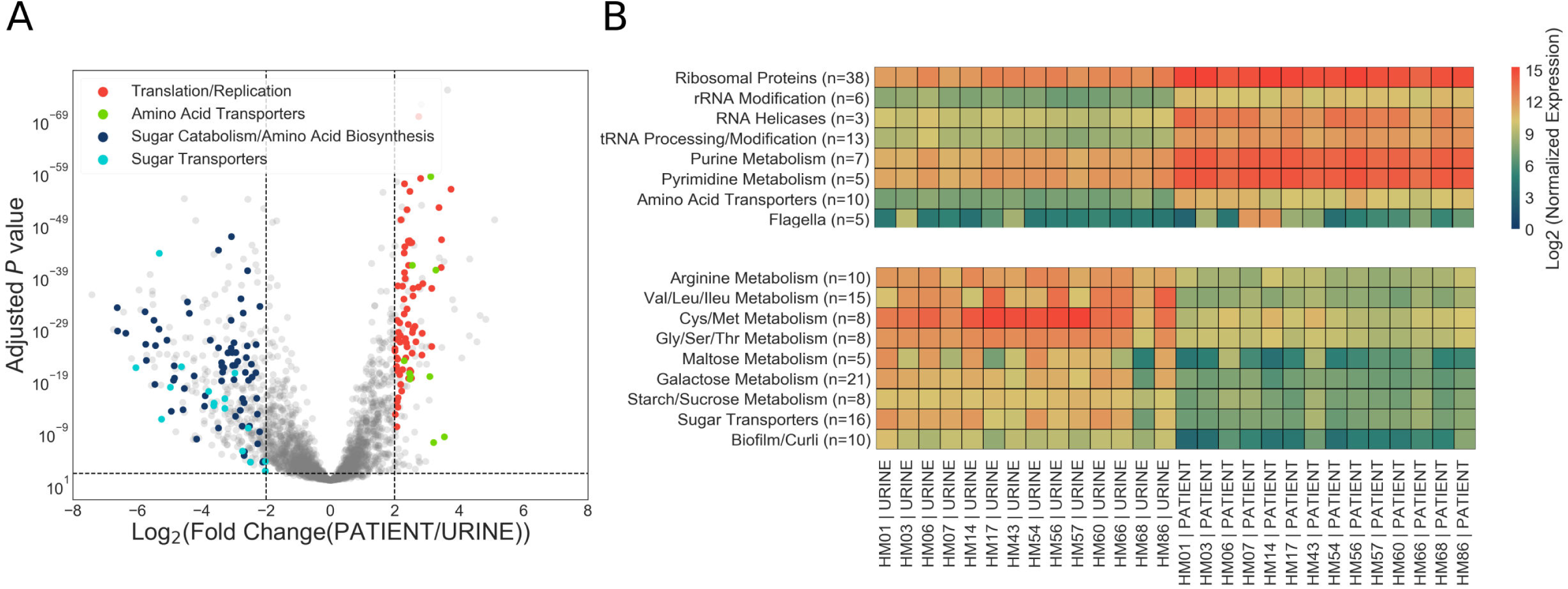
Patient-associated transcriptional signature is consistent with rapid bacterial growth. (A) The DESeq2 R package was used to compare *in vitro* urine cultures gene expression to that in patients. Each UPEC strain was considered an independent replicate (n = 14). Genes were considered up-regulated (down-regulated) if log_2_ fold change in expression was higher (lower) than 2 (vertical lines), and *P* value < 0.05 (horizontal line). Using these cutoffs, we identified 149 upregulated genes, and 343 downregulated genes. GO/pathway analysis showed that large proportion of these genes belonged to one of the 4 functional categories (see legend) (B) Mean normalized expression for genes belonging to differentially expressed functional categories/pathways. Number of up(down)-regulated genes belonging to each category is indicated next to the category name.

We also observed infection-specific downregulation of pathways involved in amino acid biosynthesis and sugar metabolism, and a general switch from expression of sugar transporters to that of amino acid transporters (Fig. 3B) (with the exception of 4 sugar transporters that were expressed at higher levels in patients: *ptsG*, *fruA*, *fruB*, and *gntU*. Fig. S6). Downregulation of sugar catabolism genes and upregulation of amino acid transporters suggest a metabolic switch to a more specific catabolic program as well as a scavenger lifestyle as examined in detail below.

### A shift in metabolic gene expression during UTI to optimize growth potential

During our analysis, we observed that 99% (on average 2621/2653 genes) of core genome was expressed during *in vitro* culture, in contrast to only 94% in patient samples (2507/2653 genes). Patient samples also contained higher proportion of genes expressed at low levels when compared to *in vitro* samples. (Fig. S5). Moreover, we noted that the majority of differentially expressed genes were downregulated in patients (343/492 differentially expressed genes). On the other hand, 30% of all upregulated genes (48/149) were ribosomal proteins. Together, these data gave us the first indication that UPEC may undergo a global gene expression reprogramming during urinary tract infection.

Bacterial growth laws postulate that bacteria dedicate a fixed amount of cellular resources to the expression of ribosomes and metabolic machinery. As a consequence, higher growth rates are achieved by allocating resources to ribosome expression at the expense of metabolic machinery production (16–21). However, this resource reallocation between ribosomal and metabolic gene expression has not yet been measured *in vivo*.

First, we wanted to determine what proportion of the total transcriptome is dedicated to core genome expression. We first hypothesized that during infection transcription could shift from the core genome to the accessory genome, which is enriched for virulence factors. However, we found that approximately 50% of total reads mapped to the core genome regardless of whether the bacteria were isolated from the patients or cultured *in vitro* (Fig. 4A). Therefore, our data indicated that a fixed proportion of cellular resources were being dedicated to expression of conserved ribosomal and metabolic machinery, regardless of external environment.

**Fig. 4.**
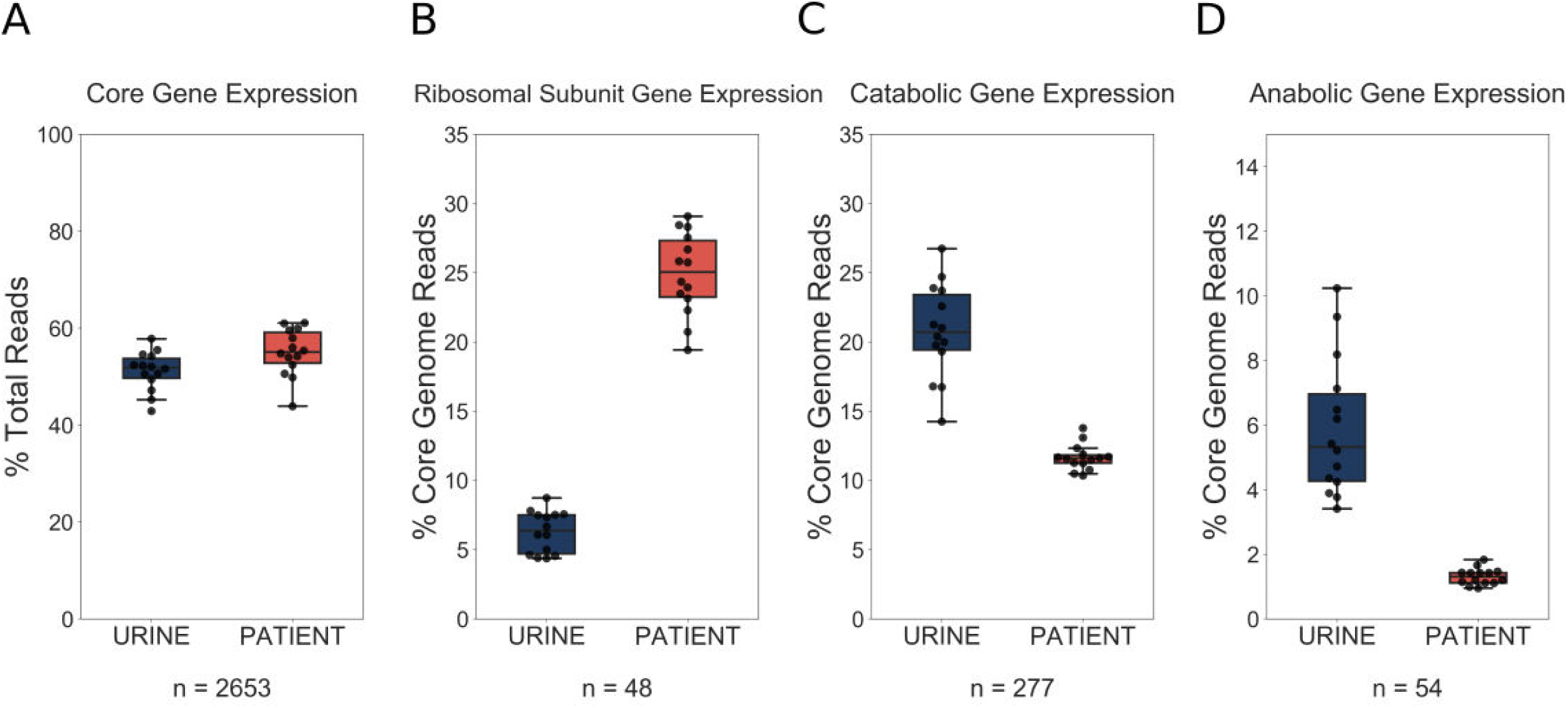
UPEC optimize growth potential via resource reallocation during UTI. (A) Percentage of reads that aligned to the core genome (2653 genes) out of total mapped reads. (B) Percentage of core genome reads that mapped to R-proteins (ribosomal subunit proteins, 48 genes). (C) Percentage of core genome reads that mapped to catabolic genes (defined as genes regulated by Crp and present in the core genome (277 genes). (D) Percentage of core genome reads that mapped to amino acid biosynthesis genes (54 genes)

We next looked at r-protein expression. Remarkably, we found that almost 25% of core genome reads mapped to r-proteins during infection, while this number was only 7% during exponential growth in urine (Fig. 4B). These findings support the idea of extremely fast UPEC growth during UTI. Furthermore, this increase in r-protein expression correlated with a marked decrease in the proportion of core genome reads dedicated to the expression of catabolic genes (20% *in vitro*, 11% in patients, Fig. 4C) and amino acid biosynthesis genes (5% *in vitro*, 1% in patients, Fig. 4D). Thus, our data highlight a dramatic and conserved resource reallocation from metabolic gene expression to replication and translational gene expression during human UTI. We postulate that this resource reallocation is required to facilitate the rapid growth rate of UPEC in the host, which has been previously documented (22).

### Increase in r-protein transcripts is an infection-specific response

Doubling time during exponential growth in urine is longer than the doubling time during exponential growth in rich media, such as LB (23). Thus, we wanted to determine whether the differences between the infection-specific and *in vitro* transcriptomes are due to longer doubling times of UPEC cultured in urine. For that purpose, one of the clinical strains, HM43, was cultured in LB, and in a new batch of filter sterilized urine. Using the optical density (OD) curves shown in Fig 5A, we estimated the doubling time of HM43 during exponential growth in LB to be approximately 33 min and the doubling time in urine to be 54 minutes. In addition, we sequenced RNA from 3-hour-old LB cultures, 3-hour-old urine cultures and from the urine of CBA/J mice, 48 hours after intraurethral inoculation with HM43.

**Fig. 5.**
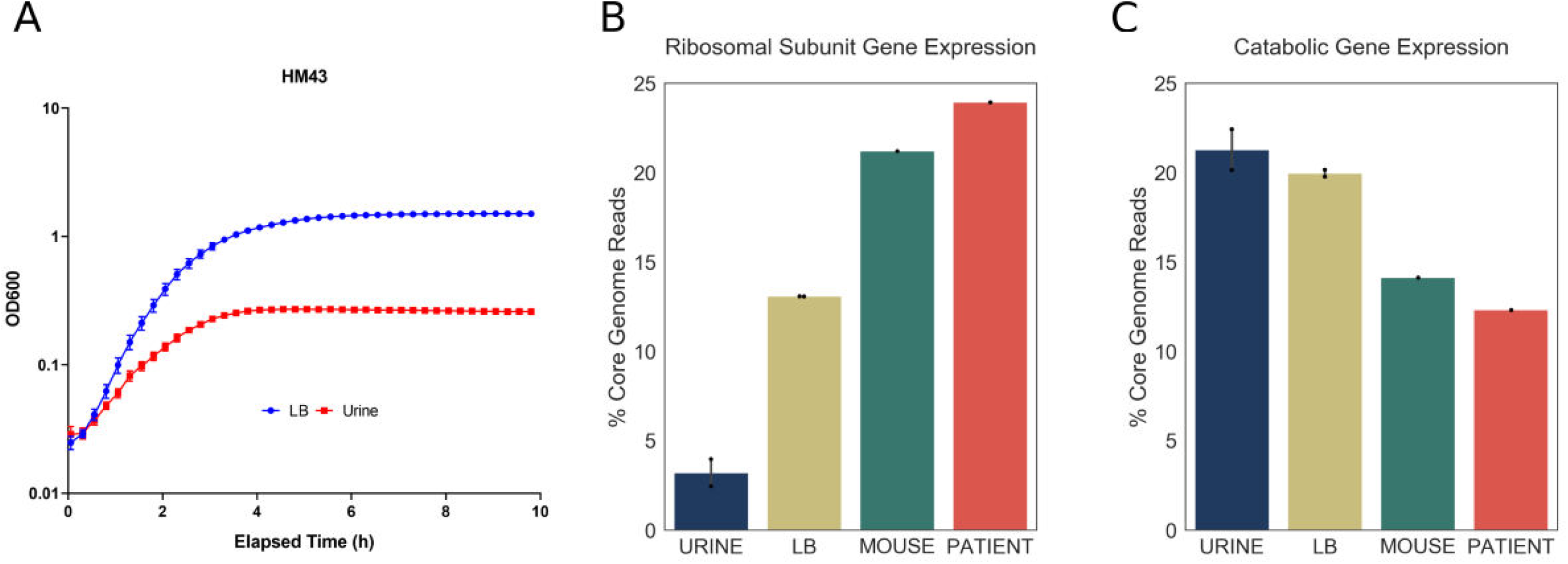
Increased expression of ribosomal subunit transcripts is a host specific response. (A) OD curve for HM43 strain cultured in LB and filter-sterilized urine. (B) Percentage of HM43 core genome reads that mapped to ribosomal subunit proteins under different conditions (URINE: *in vitro* culture in filter-sterilized urine, LB: *in vitro* culture in LB, MOUSE: mice with UTI, PATIENT: human UTI. (C) Percentage of HM43 core genome reads that mapped to catabolic genes under different conditions.

We then determined the proportion of r-protein transcripts in the HM43 transcriptomes isolated from urine and LB cultures. Consistent with our previous experiments, this proportion was very small in urine culture (4%). Interestingly, while the proportion of r-protein transcripts was approximately three times larger in LB cultures compared to urine, it was still significantly lower compared to what we observe during infection (Fig 5B). In contrast, the bacterial transcriptome during mouse infection exhibited r-protein expression that was similar to the human infection (Fig 5B). Additionally, the proportion of the transcriptome dedicated to catabolic gene expression was highest during urine cultures and lowest during mouse and human infections, indicating a negative correlation between the expression of r-protein and sugar catabolism genes. (Fig 5C). Overall, we show that exponential growth in rich medium alone cannot recapitulate the transcriptional signature observed during human infection. Taken together, our data suggest that the resource reallocation described in this study is an infection-specific response.

### Environment-responsive regulators facilitate patient-specific gene expression program

We next sought to identify potential regulators involved in resource reallocation that facilitate the infection-specific UPEC gene expression program. To do so, we performed gene set enrichment analysis (GSEA) on *E. coli* co-regulated genes (regulons). This analysis allowed us to identify regulons enriched in differentially expressed genes. We identified 22 transcriptional factors whose regulon’s expression was statistically different between infection and *in vitro* cultures (Table S3). 18/22 regulons were expressed at higher level during *in vitro* culture, and eight representative regulons are shown in Fig. 6. Overall, we found that these regulons accounted for 50% of differentially expressed genes that were determined to be significantly down-regulated. In contrast, only 6% of upregulated genes belonged to the 4 regulons that were expressed at higher levels during infection. These included genes involved in the SOS response, as well as purine synthesis (Table S3).

**Fig. 6.**
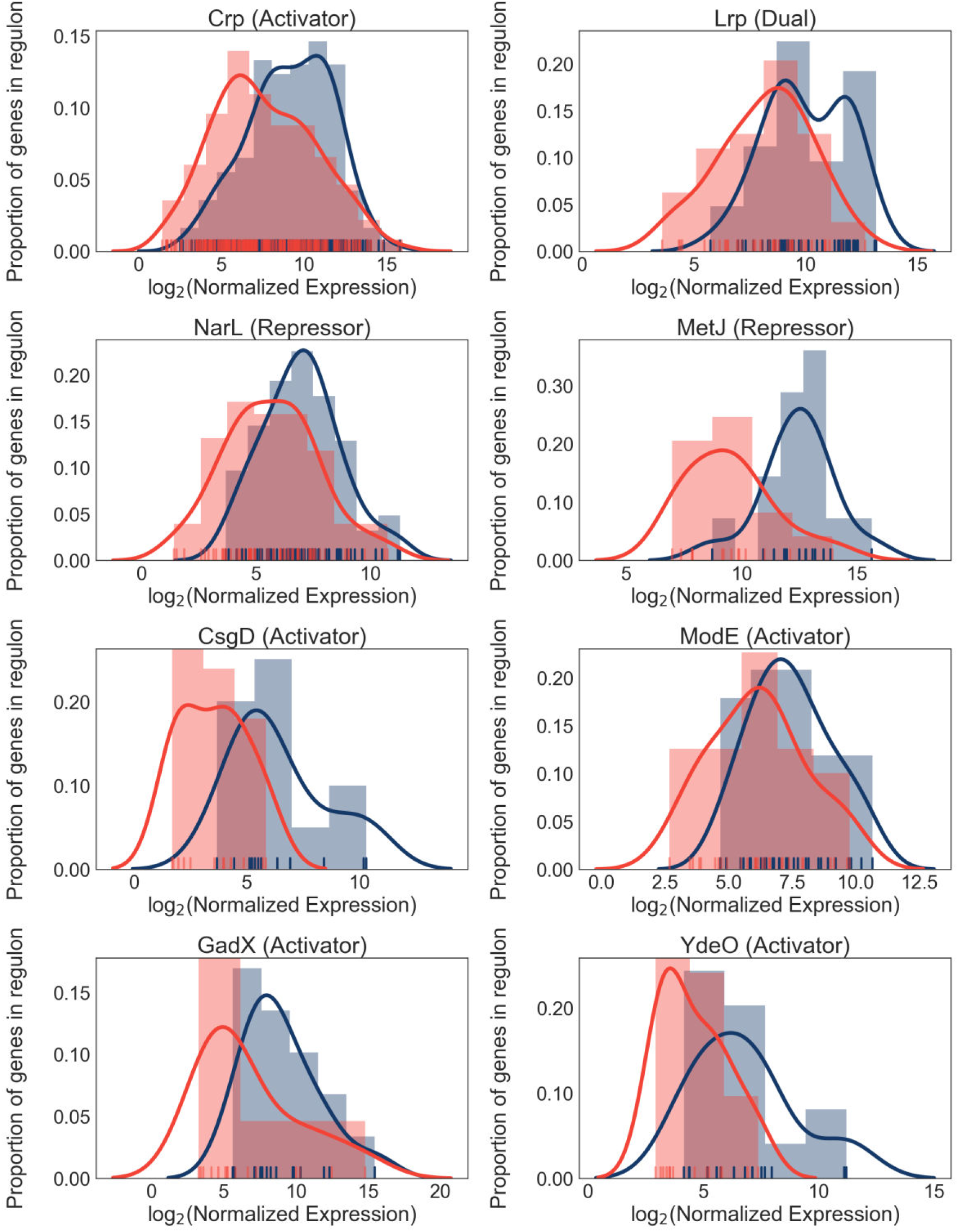
Differential regulon expression suggests role for multiple regulators in resource reallocation. Regulon expression for 8 out of 22 regulons enriched for genes downregulated in the patients. Expression of each gene in the regulon during *in vitro* culture (blue) or during UTI (red) is shown along the x-axis. Histograms show proportion of genes in the regulon expressed at any given level.

In support of our previous data, the expression of catabolic genes controlled by the Crp regulator was lower in patients compared to urine cultures. In conjunction with the previously described role for Crp in resource reallocation (21), our *in vivo* findings strongly suggest that catabolite repression plays an important role in bacterial growth rate during UTI. Interestingly, other regulators identified in this analysis (NarL, ModE, MetJ, GadE, YdeO) are known sensors of environmental cues, suggesting that the infection-specific gene expression program may be driven by additional environmental signals. Taken together, we propose a model where simultaneous sensing of multiple environmental cues in the urinary tract leads to the global down-regulation of multiple metabolic regulons during infection. The cellular resources (*e.g*., RNA polymerase) that are freed as a result are then allocated to the transcription of genes (for example, r-proteins), which are required to maintain rapid growth rate.

## Discussion

As UPEC causes one of the most prevalent bacterial infections in humans, the virulence mechanisms of UPEC infection have been well-characterized. However, while we know that these virulence strategies (*e.g.*, iron acquisition, adhesion, immune evasion) are essential for establishing infection, UPEC strains can differ dramatically in the specific factors that are utilized. Additionally, our data indicate that the expression of virulence factors can change from patient to patient, suggesting that the need for a specific factor might vary during the course of the infection.

In this study, we set out to uncover universal bacterial features during human UTIs, regardless of the stage of the infection or patient history. To do so, we performed transcriptomic analysis on bacterial RNA isolated directly from the urine of 14 patients and compared it to the gene expression of identical strains cultured to mid-exponential phase in sterile urine. Our analysis focused on the core genome as opposed to the more commonly studied accessory genome, which contains the majority of the classical virulence factors. This allowed us to identify a remarkably conserved gene expression signature shared by all 14 UPEC strains during UTI.

Although frequently overlooked, bacterial metabolism is an essential component of bacterial pathogenesis. Since the core genome is enriched for metabolic genes, we anticipated that our study, for the first time, would illuminate the UPEC metabolic state during human infection. Our data revealed an infection-specific increase in ribosomal protein expression in all 14 UPEC isolates, which was suggestive of bacteria undergoing rapid growth. While we did observe increased r-protein expression in exponentially growing UPEC cultured in LB, these transcripts were dramatically more abundant in the context of infection (human or mouse). These data suggest that UPEC are growing faster in the host in comparison to exponential growth in LB and supports recent studies that have documented very rapid UPEC growth rate in patients (22, 24).

Importantly, our analysis reveals how this growth rate can be achieved. We found that regardless of external environment, ~50% of total gene expression is allocated to the core genome, consisting of metabolic and replication machinery, which mediate bacterial growth potential. When the infection-specific transcriptome was compared to that of UPEC cultured to mid-exponential phase in urine, we observed that elevated levels of ribosomal transcripts correlated with decreased levels of metabolic gene expression. We propose that this reallocation of resources within the core genome drives the rapid growth rate of UPEC during infection.

This resource reallocation is equivalent to what has been described as the bacterial ‘growth law’. Based on *in vitro* studies, the growth law proposes that increases in ribosomal gene expression occurs at the expense of a cell’s metabolic gene expression (17, 19). Our analysis of UPEC gene expression directly from patients is consistent with this hypothesis. In addition, regulatory network analysis revealed that multiple metabolic regulons exhibit decreased transcript levels in patients suggesting an actively regulated process. In contrast, synthesis of ribosomal RNA (rRNA) coordinates the expression of ribosomal proteins by a translational feedback regulation mechanism (25, 26). rRNA synthesis is proposed to be regulated by the competition of RNA polymerase between transcription of rRNA operons and that of other genes, with some studies suggesting that mid-log growing cells might require almost all RNA polymerase dedicated to rRNA synthesis (30–33). Thus, decreased metabolic gene expression could allow the cell to shift its allocation of RNA polymerase towards rRNA synthesis and as a result, ribosomal protein expression. Although we cannot exclude other mechanisms, we propose that the reallocation of RNA polymerase molecules from metabolic genes to rRNA and ribosomal protein genes is a common feature adopted by diverse UPEC to promote rapid growth during UTI.

Three recent studies have attempted to characterize UPEC gene expression in patients with UTIs (15, 27, 28). These studies focused on the importance of virulence factor expression in specific strains and have demonstrated changes in gene expression between infection and *in vitro* cultures. It should be noted that all of these studies, as well as our own, were performed using bacterial RNA isolated from patient urine (that was immediately stabilized upon collection). As a result, we cannot exclude the possibility that gene expression of UPEC residing in the bladder may differ from UPEC isolated from patient urine. However, the fact remains that 14 patients with different histories of UTIs all harbored a population of actively dividing bacteria in a remarkably specific metabolic state, which we have also recapitulated in a mouse model of infection.

These findings raise a number of interesting questions. Firstly, how is rapid growth rate beneficial to UPEC and how does it influence the tempo and mode of bacterial evolution, especially with regards to genomic integrity and the acquisition of antibiotic resistance? Secondly, what are the external cues that launch the infection-specific transcriptional response? While our study was not designed to identify infection-specific metabolites, our regulatory network analysis suggests that multiple environmental cues might reinforce the suppression of metabolic gene expression. We suggest that identifying and targeting these environmental cues is a promising approach to limit UPEC growth during UTI and gain the upper hand on this pathogen.

## Methods

### Study design

Sample collection was previously described (15). Briefly, a total of 86 female participants, presenting with symptoms of lower UTI at the University of Michigan Health Service Clinic in Ann Arbor, MI in 2012, were enrolled in this study. The participants were compensated with a $10 gift card to a popular retail store. Clean catch midstream urine samples from participants were immediately stabilized with two volumes of RNAprotect (Qiagen) to preserve the *in vivo* transcriptional profile. De-identified patient samples were assigned unique sample numbers and used in this study. Of the 86 participants, 38 were diagnosed with UPEC-associated UTIs (15). Of these, 19 samples gave us sufficient RNA yield of satisfactory quality. Five were used for a pilot project (15), the remaining 14 were used in this study.

### Genome sequencing and assembly

The genomic DNA from clinical strains of *E. coli* were isolated with CTAB/phenol-chloroform based protocol. Library preparation and sequencing were performed on PacBio RS system at University of Michigan Sequencing Core. *De novo* assemblies were performed with <u>canu</u> *de novo* assembler (29) with all the parameters set to default mode and correction phase turned on. We used progressiveMauve (30) with default parameters to align draft assemblies of clinical strains, and MG1655. Maximum likelihood tree was generated based on core SNPS modeled with a general-time reversible (GTR) model in RAxML (31).

### Phylogroup analysis

Phylogroups were assigned using an in-house script based on the presence and absence of primer target sequences and typing scheme (32).

### Bacterial culture conditions

Human urine was pooled from four age-matched healthy female volunteers. Overnight cultures of clinical isolates were washed once in human urine, then 250 *μ*l of overnight culture was added to 25 ml of filter-sterilized human urine and cultured statically at 37C for 2 hours. Six milliliters of this culture were stabilized with RNAprotect (Qiagen) and used for RNA purification.

### Antibiotic resistance profiling

As described in (15), identity and antibiotic resistance profiles of UPEC isolates were determined using a VITEK2 system (BioMerieux).

### RNA isolation and sequencing

RNA isolation protocol was previously described (15). Briefly, samples were treated with proteinase K and total RNA was isolated using Qiagen RNAeasy minikit. Turbo DNase kit (Ambion) was used to remove contaminating DNA. Bacterial content of patient samples was enriched using MICROBenrich kit (Ambion). Library preparation and sequencing was performed by University of Michigan sequencing core. Illumina ScriptSeq v2 library kit was used to construct rRNA-depleted stranded cDNA libraries. These were sequenced using Illumina HiSeq2500 (single end, 50-bp read length).

### Characterization of virulence factors’ gene expression

We compiled a literature search-based list of virulence factors belonging to different functional groups. Sequences for each virulence factor gene were extracted from reference UPEC genomes (either CFT073 or UTI89). Presence or absence of each virulence factor within clinical genomes was determined using BLAST (with percent identity ≥ 80% and percent coverage ≥ 90%, e-value ≤ 10^−6^). Hierarchical clustering of strains based on presence or absence of virulence factors was performed using Python’s scipy.cluster.hierarchy.linkage function with default parameters. Heatmaps of virulence factors’ gene expression in urine and in patients show normalized transcripts per million (TPMs) (same as for correlation analysis and PCA, see below).

### RNAseq Data Processing

A custom bioinformatics pipeline was used for the analysis (github.com/ASintsova/rnaseq_analysis). Raw fastq files were processed with Trimmomatic (33) to remove adapter sequences and analyzed with FastQC to assess sequencing quality. Mapping was done with bowtie2 aligner (34) using default parameters. Alignment details can be found in Table S1. Read counts were calculated using HTseq htseq-count (union mode) (35).

### Quality control

Since some of our clinical samples yielded lower numbers of bacterial reads than desired (Table S1), we performed a comprehensive quality assurance to determine if the sequencing depth of our clinical samples was sufficient for our analysis (see Saturation curves and Gene expression ranges analysis below, Fig. S4, Fig. S5). Overall, all patient samples except for HM66 passed quality control (see gene expression ranges analysis, Fig. S5). While we elected to keep all of the strains in our subsequent analysis, this observation explains why the patient HM66 sample appears as an outlier in Fig. 2.

### Saturation curves

We created saturation curves for each of our sequencing files to assess whether we have sufficient sequencing depth for downstream analysis. Each sequencing file was subsampled to various degrees and number of genes detected in those subsamples (y-axis) was graphed against number of reads in the subsample (x-axis). As expected, all of the *in vitro* samples reached saturation (Fig S4, blue lines). Unfortunately, 6 out of our 14 samples did not reach saturation, which warranted us to investigate further (see Gene expression ranges analysis) Fig S4, red lines).

### Core genome identification

Core genome for 14 clinical isolates and MG1655 was determined using get_homologues (36). We explored multiple parameter values for our analysis and their effect on final core genome, in the end we set the cut off of 90% of sequence identity and 50% sequence coverage (similar results were obtained when using different cutoffs). The intersection of three algorithms employed by get_homologues contained 2653 gene clusters.

### Gene expression ranges analysis

Due to low sequencing depth of 6 of our isolates, we were worried we would not be able to detect genes expressed at low levels in those samples. To evaluate whether we were losing information about low-level expression, we compared a number of genes in the core genome that were expressed at different levels (1000 TPMS, 100 TPMS, 10 TPMS and 1 TPM) between clinical samples that reached saturation (Fig. S5A) and those that did not (Fig. S5B). Only one of the clinical samples (HM66) seemed to lack genes expressed in the range of 1-10 TPMs. Thus, we conclude that all but one sample (HM66) had sufficient coverage for downstream analysis.

### Pearson correlation coefficient calculation and PCA analysis

For PCA and correlation analysis, transcript per million (TPM) was calculated for each gene, TPM distribution was then normalized using inverse rank transformation. Pearson correlation and PCA was performed using python Python sklearn library. Jupyter notebooks used to generate the figures are available at https://github.com/ASintsova/HUTI_RNAseq

### Differential expression analysis

Differential expression analysis was performed using DESeq2 R package (37). Genes with log2 fold change of greater than 2 or less than −2 and adjusted *p* value (Benjamini-Hochberg adjustment) of less than 0.05 were considered to be differentially expressed. DESeq2 normalized counts were used to generate Fig. 3 and 6. Pathway analysis was performed using R package topGO (38).

### RNA sequencing of HM43 from mouse model of UTI

Forty CBA/J mice were infected using the ascending model of UTI as previously described (39, 40). Briefly, 40 six-week-old female mice were transurethrally inoculated with 10^8^ CFU of UPEC isolate HM43. 48 hours post infection urine was collected from each mouse directly into bacterial RNAprotect (Qiagen). All collected urine was pooled together and pelleted, and immediately placed in the −80°C freezer. This collection was repeated every 45 minutes five more times, resulting in six collected pellets consisting of bacteria and eukaryotic cells.

For *in vitro* controls, UPEC strain HM43 was cultured overnight in LB. The next morning, the culture was spun down, and the pellet washed twice with PBS. LB or pooled human urine was then inoculated with the washed bacteria at a ratio of 1:100 and incubated with shaking at 37°C for 3 hours. Cultures were then treated with bacterial RNAprotect (Qiagen), pellets collected and stored at −80°C.

All the pellets were treated with both lysozyme and proteinase K, and then total RNA was extracted using RNAeasy kit (Qiagen). Genomic DNA was removed using the Turbo DNA free kit (ThermoFisher). Eukaryotic mRNA was depleted using dynabeads covalently linked with oligo dT (ThermoFisher). The supernatant was collected from this treatment, and the RNA was concentrated and re-purified using RNA Clean & Concentrator kit (Zymo). Library preparation and sequencing was done as described above.

### Estimation of HM43 doubling time

For both LB and urine OD curves were performed using Bioscreen-C Automated Growth Curve Analysis System (Growth Curves USA) 8 separate times. For each time point, the mean values of the 8 replicates were used for doubling time estimation. The equation bellow was used to estimate doubling time during logarithmic growth in LB or urine, where DT is doubling time, C2 is final OD, C1 is initial OD, and ∆*t* is time elapsed between when C2 and C1 were taken.

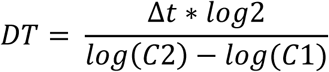

DT was calculated for every two measurements taken between 30 and 180 minutes and mean of these values is reported.

### Regulon analysis

Regulon gene sets were extracted from RegulonDB 9.4 (41) using custom Python scripts (available https://github.com/ASintsova/HUTI_RNAseq). Gene set enrichment analysis was performed using Python GSEAPY library.

### Data access

Jupyter notebooks as well as count data used to generate all the figures in this paper are available on github: https://github.com/ASintsova/HUTI_RNAseq

---

**Figure S1.**
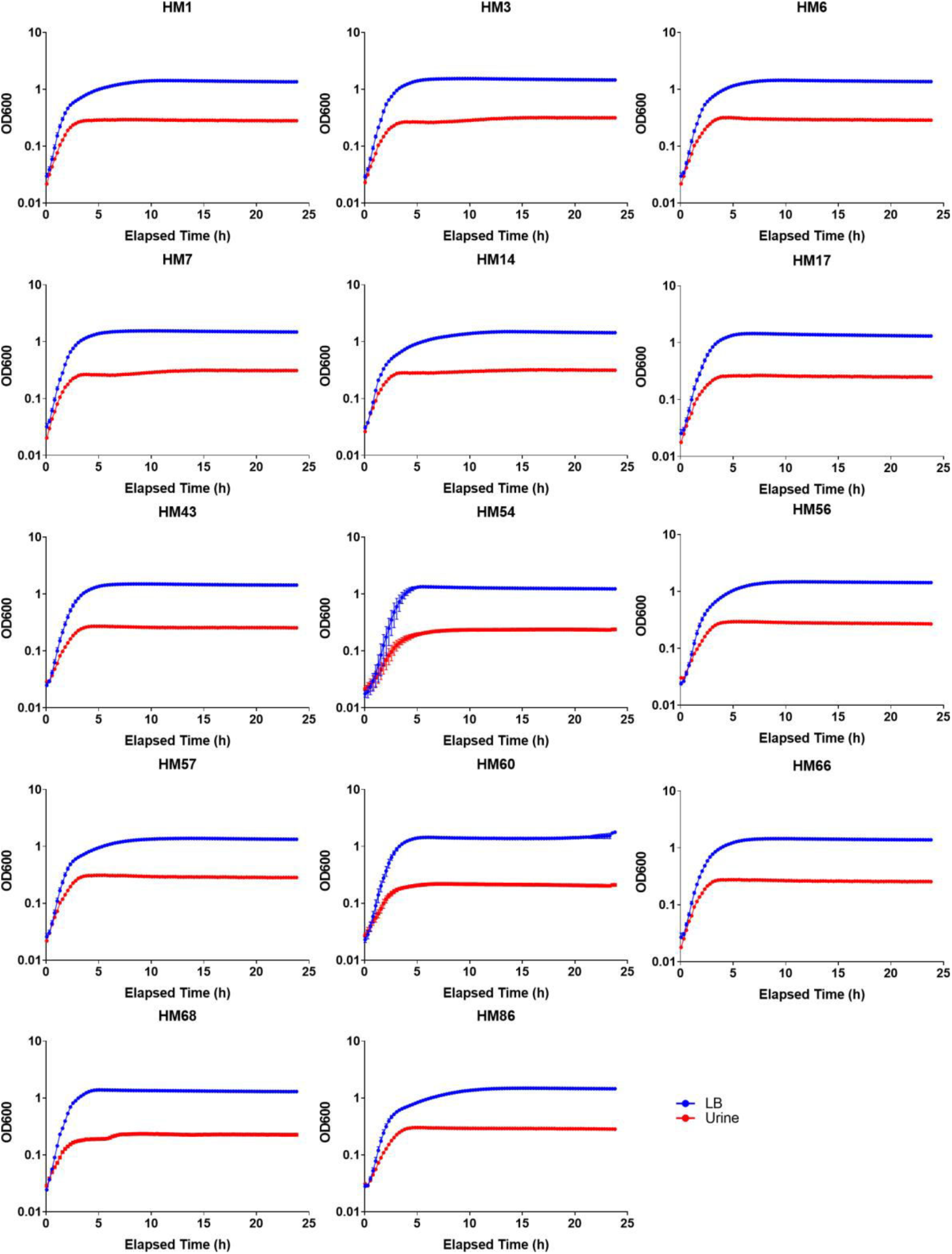
OD curves for 14 clinical UPEC strains cultured in LB or filter-sterilized urine.

**Figure S2.**
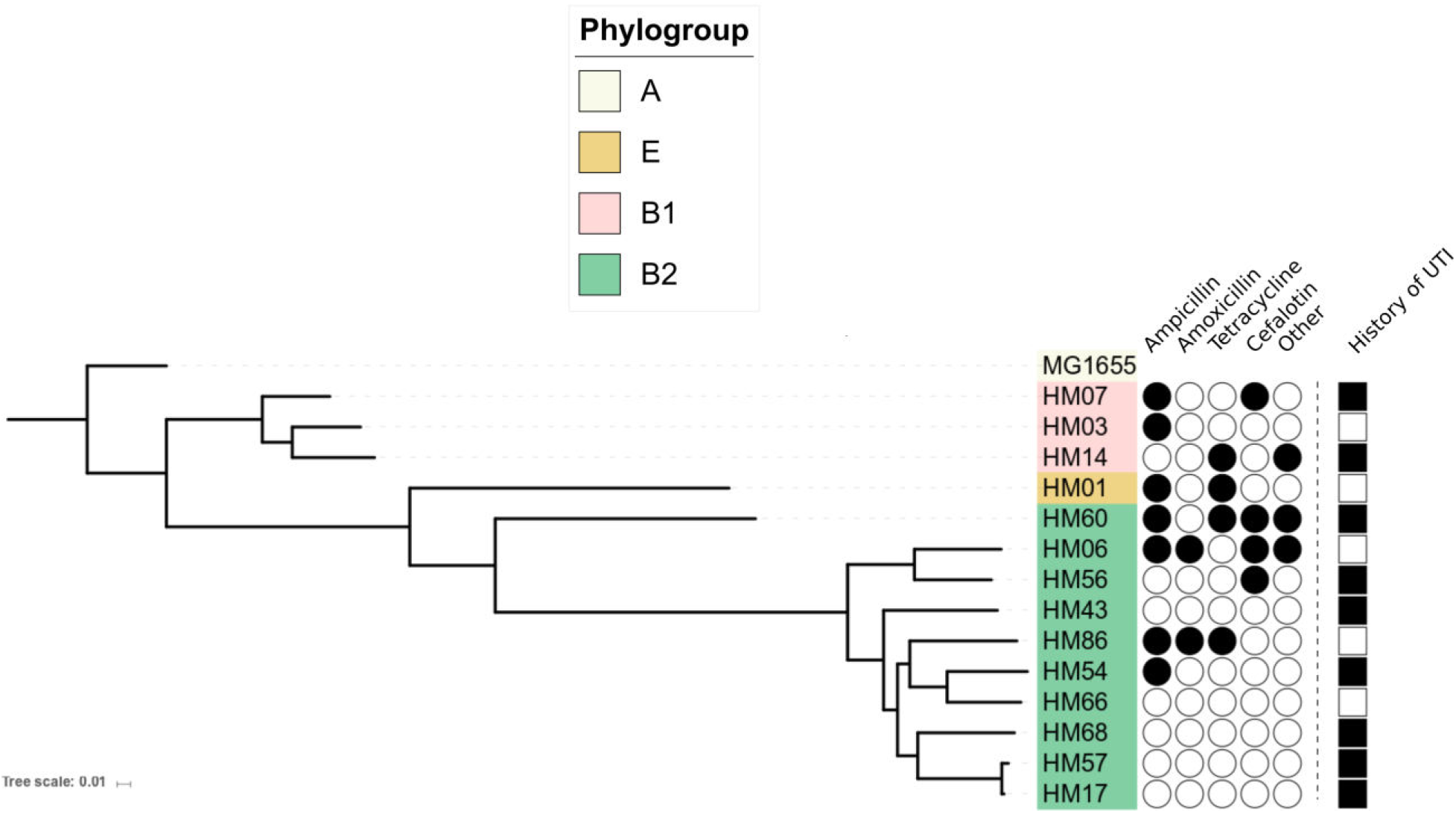
ML Phylogenetic tree reconstruction of 14 clinical UPEC strains isolated in this study using core SNPs. Antibiotic resistance profiles are indicated by filled in black circles (as determined by VITEK2 system (BioMerieux).) Patients with recurrent UTIs are indicated by filled in black square.

**Fig. S3.**
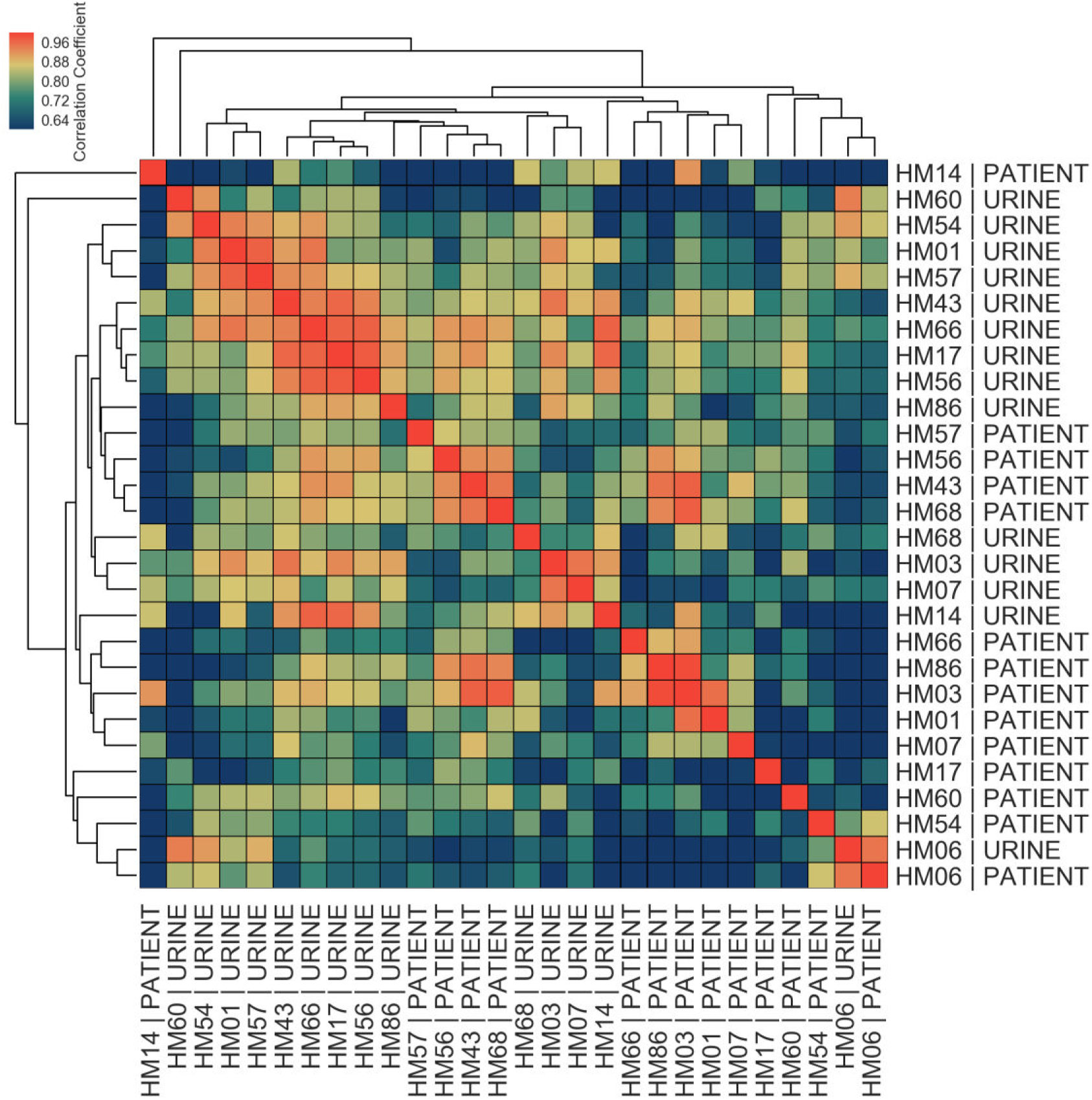
Correlations among *in vitro* and patient samples measured by Pearson correlation coefficient of normalized gene expression of 40 virulence factors plotted according to hierarchical clustering of samples.

**Figure S4.**
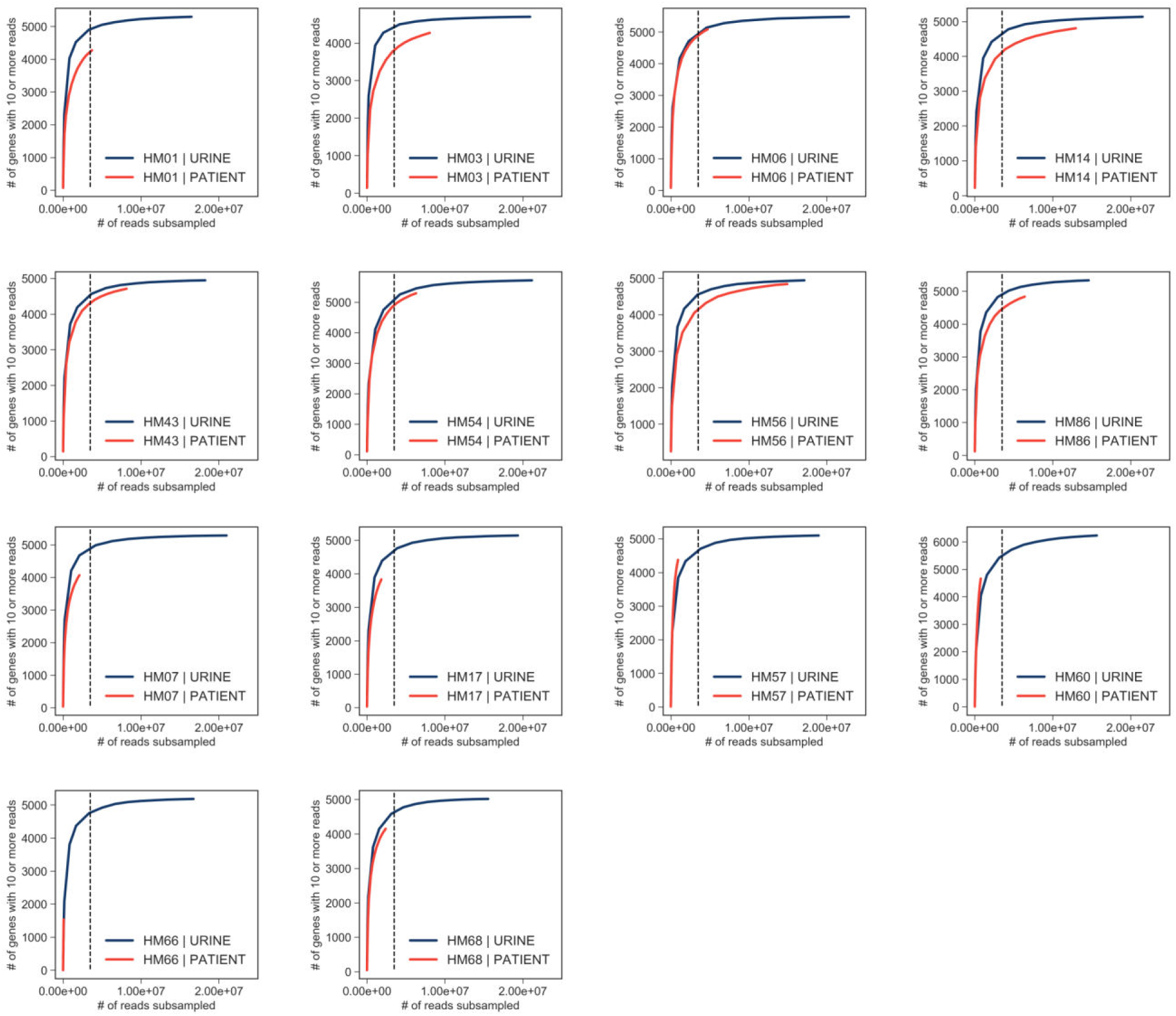
Saturation curves. Number of mapped reads was plotted against number of expressed genes detected for each sample (*in vitro* samples are shown in blue, patient samples are shown in red). Vertical line shows 3 million reads cut off at which samples appear to reach saturation.

**Figure S5.**
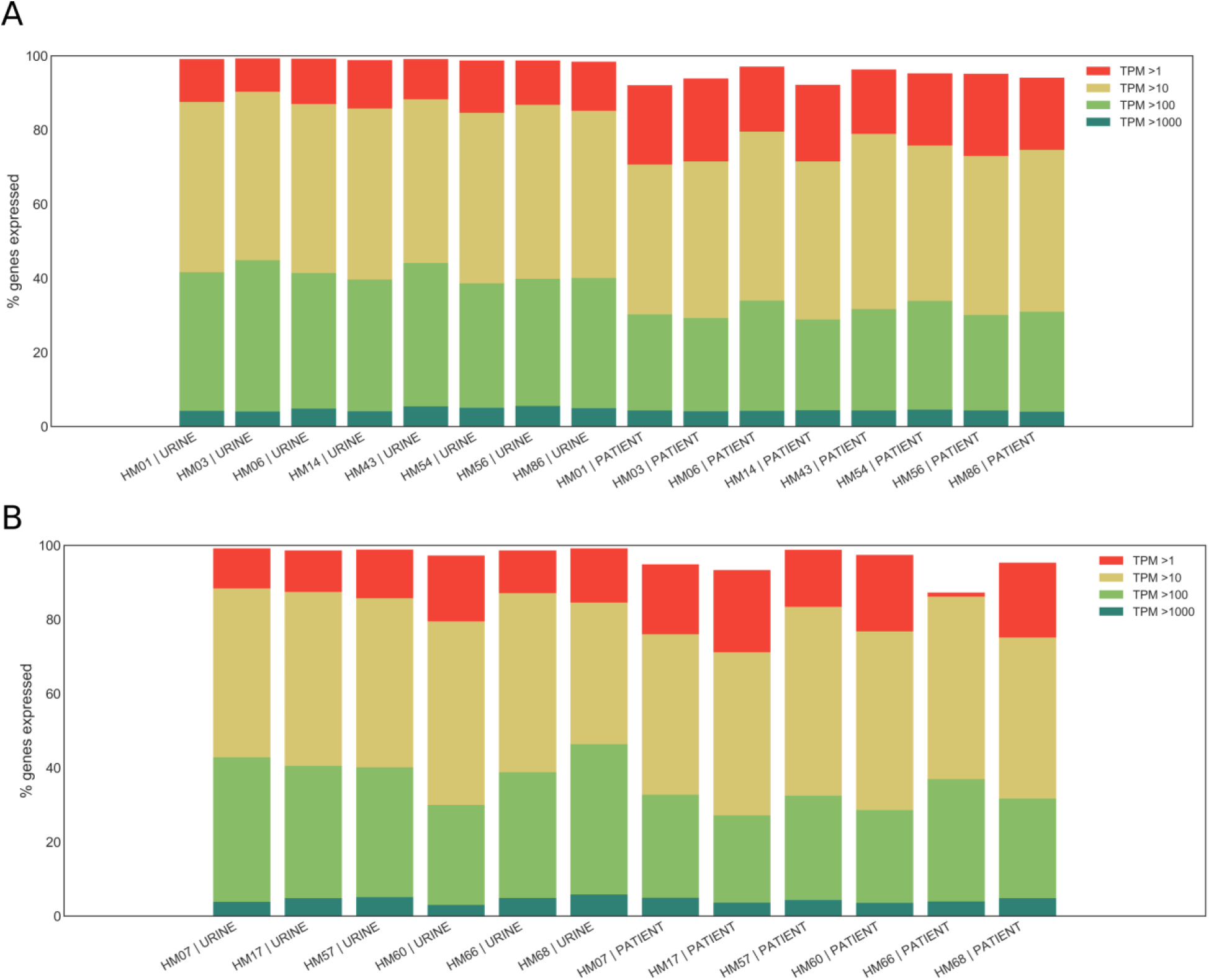
Expression ranges of core genome genes. (A) Percentage of genes in the core genome that are expressed at a given level (>1 TPM, >10 TPMs, > 100 TPMs, > 1000 TPMs, where TPMs are transcripts per million) is shown for patient samples that reached saturation (see Supplementary Figure 2) and corresponding *in vitro* samples. (B) Percentage of genes in the core genome that are expressed at a given level (>1 TPM, >10 TPMs, > 100 TPMs, > 1000 TPMs) is shown for patient samples that did not reach saturation and corresponding *in vitro* samples.

**Figure S6.**
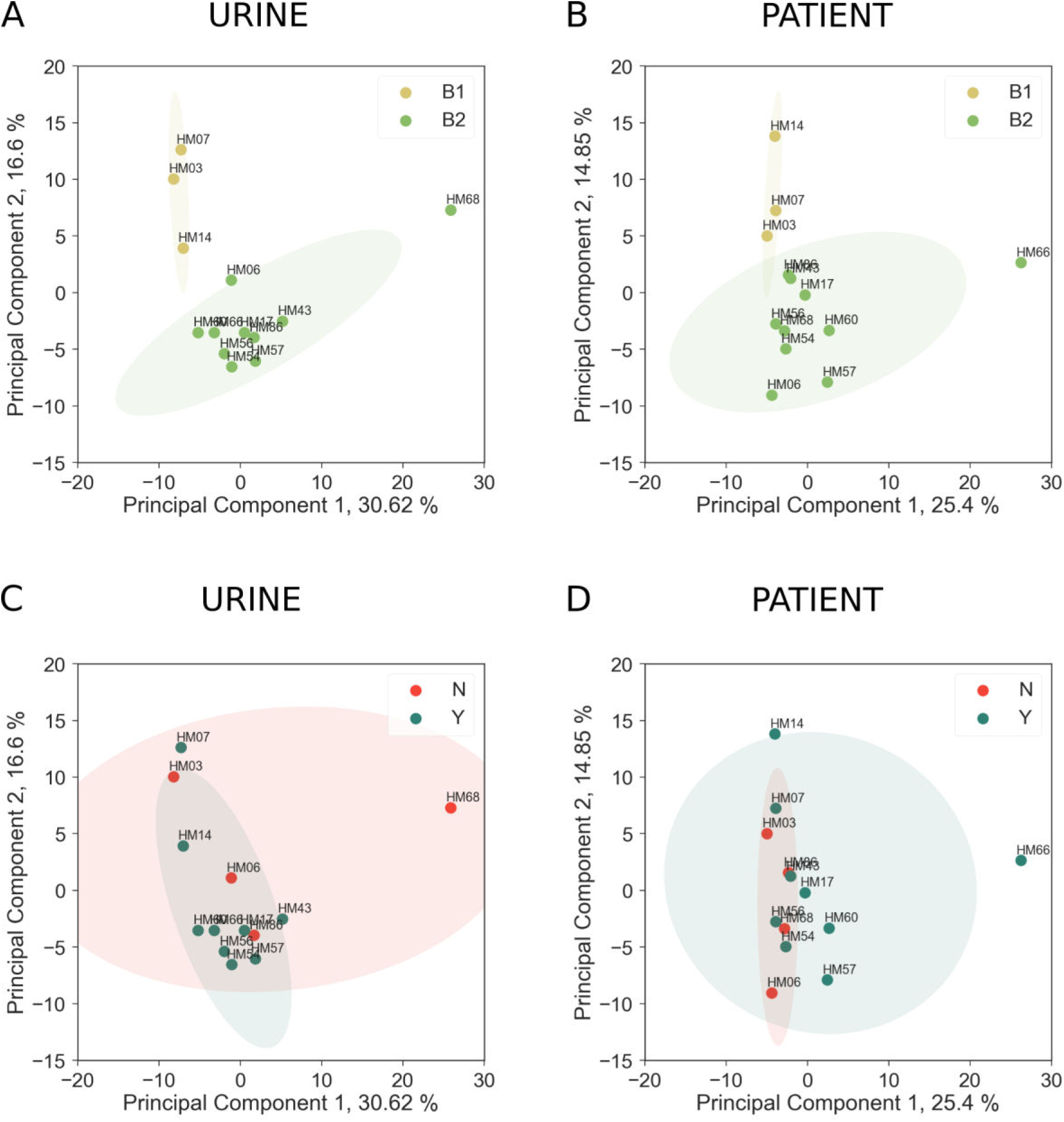
Effect of phylogenetic group on core genome expression. (A) and (C) Clustering of UPEC strains cultured in filter-sterilized urine based on PCA analysis of core genome gene expression. (B) and (D) Clustering of UPEC isolated from patients based on PCA analysis of core genome gene expression. Samples in (A) and (B) are colored based on their phylogroup designation. Samples in (C) and (D) are colored based on whether the strain was isolated from a patient with recurrent UTI (Y) or without recurrent UTI (N).

**Figure S7.**
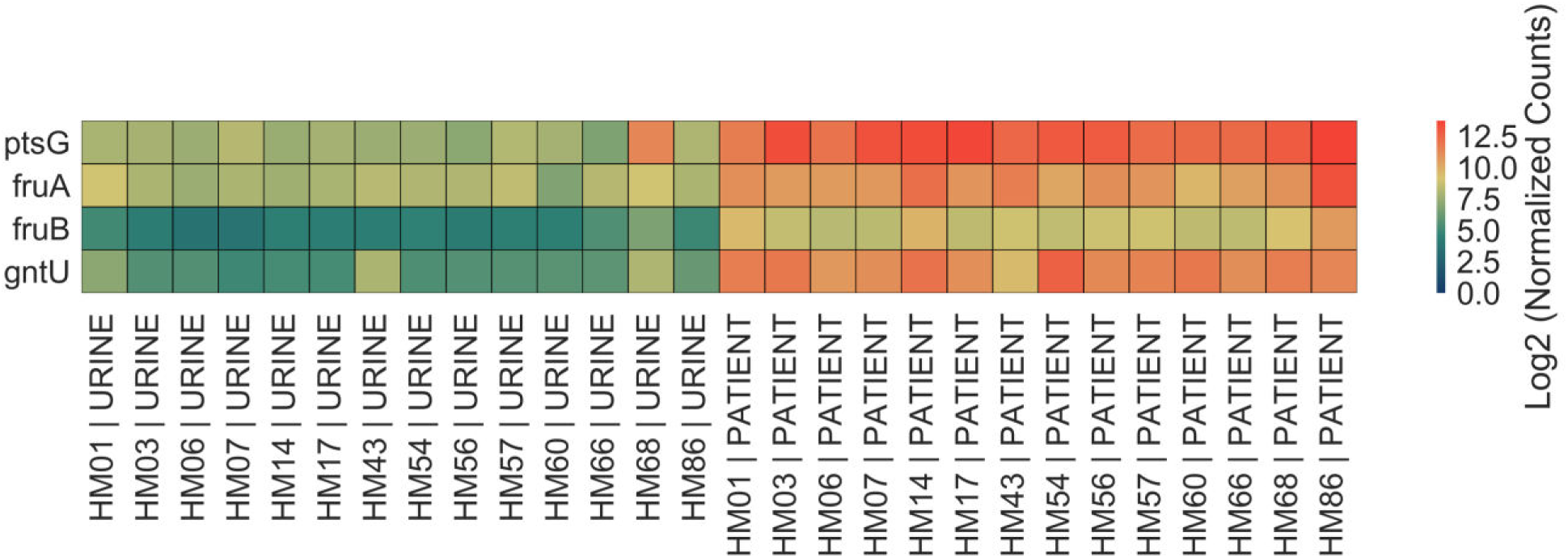
Gene expression of four sugar transporters upregulated in UTI patients. Heatmap shows Log2 of normalized gene expression of *ptsG*, *fruA*, *fruB* and *gntU* for each *in vitro* and patient sample.

**Table S1.** Summary of alignment statistics.

| **Sample:** | **Total Reads:** | **Mapped Reads:** | **% Mapped:** |
| --- | --- | --- | --- |
| **HM01 \| URINE** | 17288419 | 16480326 | 95.3 |
| **HM01 \| PATIENT** | 18496607 | 3717040 | 20.1 |
| **HM03 \| URINE** | 21354719 | 20927541 | 98 |
| **HM03 \| PATIENT** | 16544044 | 8059076 | 48.7 |
| **HM06 \| URINE** | 23359847 | 22847374 | 97.8 |
| **HM06 \| PATIENT** | 57993519 | 4709092 | 8.1 |
| **HM07 \| URINE** | 21312224 | 20980473 | 98.4 |
| **HM07 \| PATIENT** | 70804688 | 2097350 | 3 |
| **HM14 \| URINE** | 21927302 | 21533817 | 98.2 |
| **HM14 \| PATIENT** | 15944762 | 12968218 | 81.3 |
| **HM17 \| URINE** | 19790215 | 19360294 | 97.8 |
| **HM17 \| PATIENT** | 23874585 | 1842583 | 7.7 |
| **HM43 \| URINE** | 18541484 | 18239826 | 98.4 |
| **HM43 \| PATIENT** | 58306859 | 8138559 | 14 |
| **HM54 \| URINE** | 21612581 | 21162544 | 97.9 |
| **HM54 \| PATIENT** | 18000843 | 6301998 | 35 |
| **HM56 \| URINE** | 17494135 | 17130847 | 97.9 |
| **HM56 \| PATIENT** | 25408755 | 14935948 | 58.8 |
| **HM57 \| URINE** | 19253078 | 18966748 | 98.5 |
| **HM57 \| PATIENT** | 105629816 | 926795 | 0.9 |
| **HM60 \| URINE** | 15898045 | 15651916 | 98.5 |
| **HM60 \| PATIENT** | 76149837 | 764255 | 1 |
| **HM66 \| URINE** | 17184018 | 16736066 | 97.4 |
| **HM66 \| PATIENT** | 25954183 | 79859 | 0.3 |
| **HM68 \| URINE** | 15841639 | 15562711 | 98.2 |
| **HM68 \| PATIENT** | 65413931 | 2401089 | 3.7 |
| **HM86 \| URINE** | 15019669 | 14606346 | 97.2 |
| **HM86 \| PATIENT** | 10667404 | 6413794 | 60.1 |

**Supplementary Table S2:**
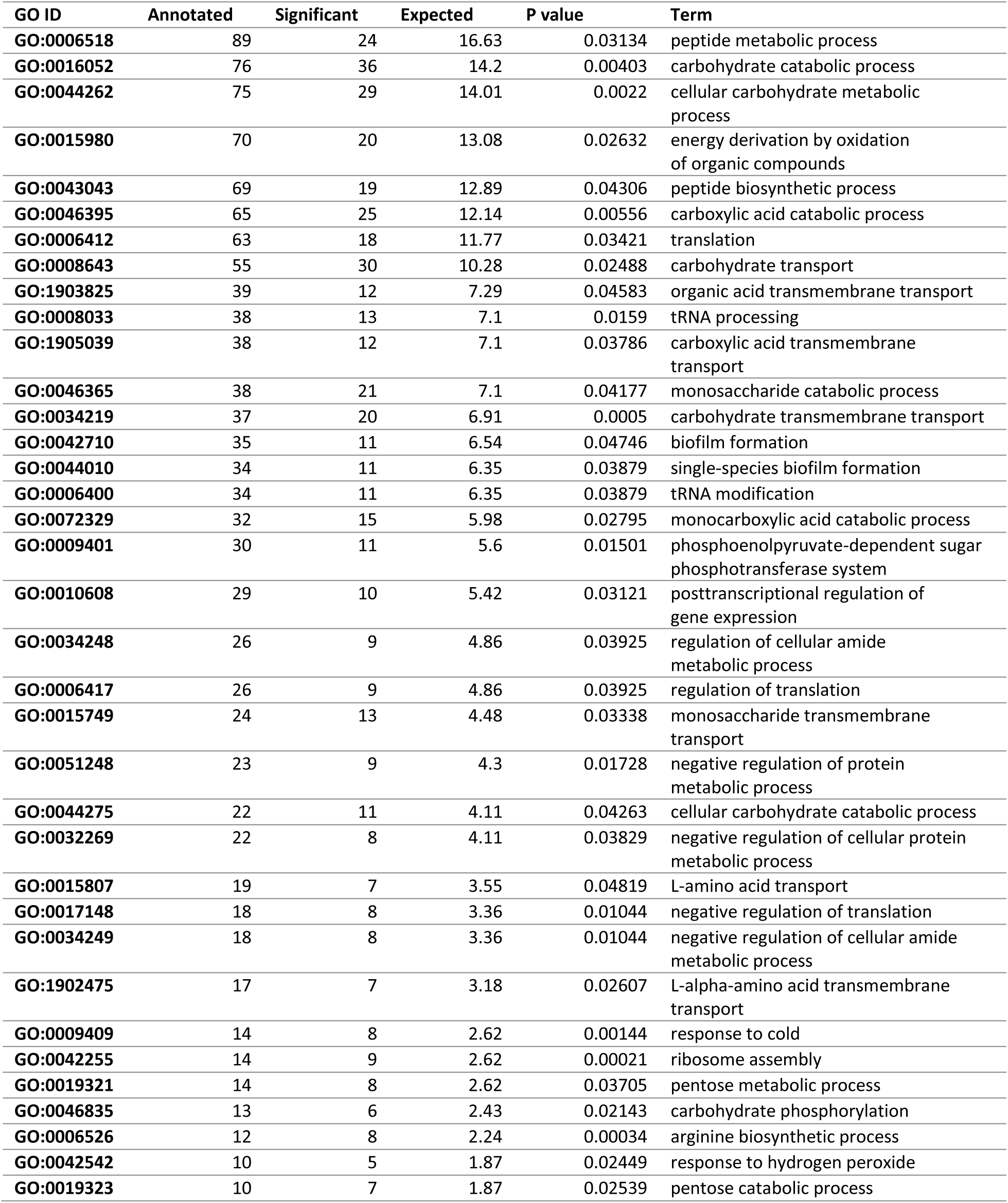
GO modules differentially expressed in UTI patients.

| GO ID | Annotated | Significant | Expected | P value | Term |
| --- | --- | --- | --- | --- | --- |
| GO:0006518 | 89 | 24 | 16.63 | 0.03134 | peptide metabolic process |
| GO:0016052 | 76 | 36 | 14.2 | 0.00403 | carbohydrate catabolic process |
| GO:0044262 | 75 | 29 | 14.01 | 0.0022 | cellular carbohydrate metabolic process |
| GO:0015980 | 70 | 20 | 13.08 | 0.02632 | energy derivation by oxidation of organic compounds |
| GO:0043043 | 69 | 19 | 12.89 | 0.04306 | peptide biosynthetic process |
| GO:0046395 | 65 | 25 | 12.14 | 0.00556 | carboxylic acid catabolic process |
| GO:0006412 | 63 | 18 | 11.77 | 0.03421 | translation |
| GO:0008643 | 55 | 30 | 10.28 | 0.02488 | carbohydrate transport |
| GO:1903825 | 39 | 12 | 7.29 | 0.04583 | organic acid transmembrane transport |
| GO:0008033 | 38 | 13 | 7.1 | 0.0159 | tRNA processing |
| GO:1905039 | 38 | 12 | 7.1 | 0.03786 | carboxylic acid transmembrane transport |
| GO:0046365 | 38 | 21 | 7.1 | 0.04177 | monosaccharide catabolic process |
| GO:0034219 | 37 | 20 | 6.91 | 0.0005 | carbohydrate transmembrane transport |
| GO:0042710 | 35 | 11 | 6.54 | 0.04746 | biofilm formation |
| GO:0044010 | 34 | 11 | 6.35 | 0.03879 | single-species biofilm formation |
| GO:0006400 | 34 | 11 | 6.35 | 0.03879 | tRNA modification |
| GO:0072329 | 32 | 15 | 5.98 | 0.02795 | monocarboxylic acid catabolic process |
| GO:0009401 | 30 | 11 | 5.6 | 0.01501 | phosphoenolpyruvate-dependent sugar phosphotransferase system |
| GO:0010608 | 29 | 10 | 5.42 | 0.03121 | posttranscriptional regulation of gene expression |
| GO:0034248 | 26 | 9 | 4.86 | 0.03925 | regulation of cellular amide metabolic process |
| GO:0006417 | 26 | 9 | 4.86 | 0.03925 | regulation of translation |
| GO:0015749 | 24 | 13 | 4.48 | 0.03338 | monosaccharide transmembrane transport |
| GO:0051248 | 23 | 9 | 4.3 | 0.01728 | negative regulation of protein metabolic process |
| GO:0044275 | 22 | 11 | 4.11 | 0.04263 | cellular carbohydrate catabolic process |
| GO:0032269 | 22 | 8 | 4.11 | 0.03829 | negative regulation of cellular protein metabolic process |
| GO:0015807 | 19 | 7 | 3.55 | 0.04819 | L-amino acid transport |
| GO:0017148 | 18 | 8 | 3.36 | 0.01044 | negative regulation of translation |
| GO:0034249 | 18 | 8 | 3.36 | 0.01044 | negative regulation of cellular amide metabolic process |
| GO:1902475 | 17 | 7 | 3.18 | 0.02607 | L-alpha-amino acid transmembrane transport |
| GO:0009409 | 14 | 8 | 2.62 | 0.00144 | response to cold |
| GO:0042255 | 14 | 9 | 2.62 | 0.00021 | ribosome assembly |
| GO:0019321 | 14 | 8 | 2.62 | 0.03705 | pentose metabolic process |
| GO:0046835 | 13 | 6 | 2.43 | 0.02143 | carbohydrate phosphorylation |
| GO:0006526 | 12 | 8 | 2.24 | 0.00034 | arginine biosynthetic process |
| GO:0042542 | 10 | 5 | 1.87 | 0.02449 | response to hydrogen peroxide |
| GO:0019323 | 10 | 7 | 1.87 | 0.02539 | pentose catabolic process |

**Table S3:**
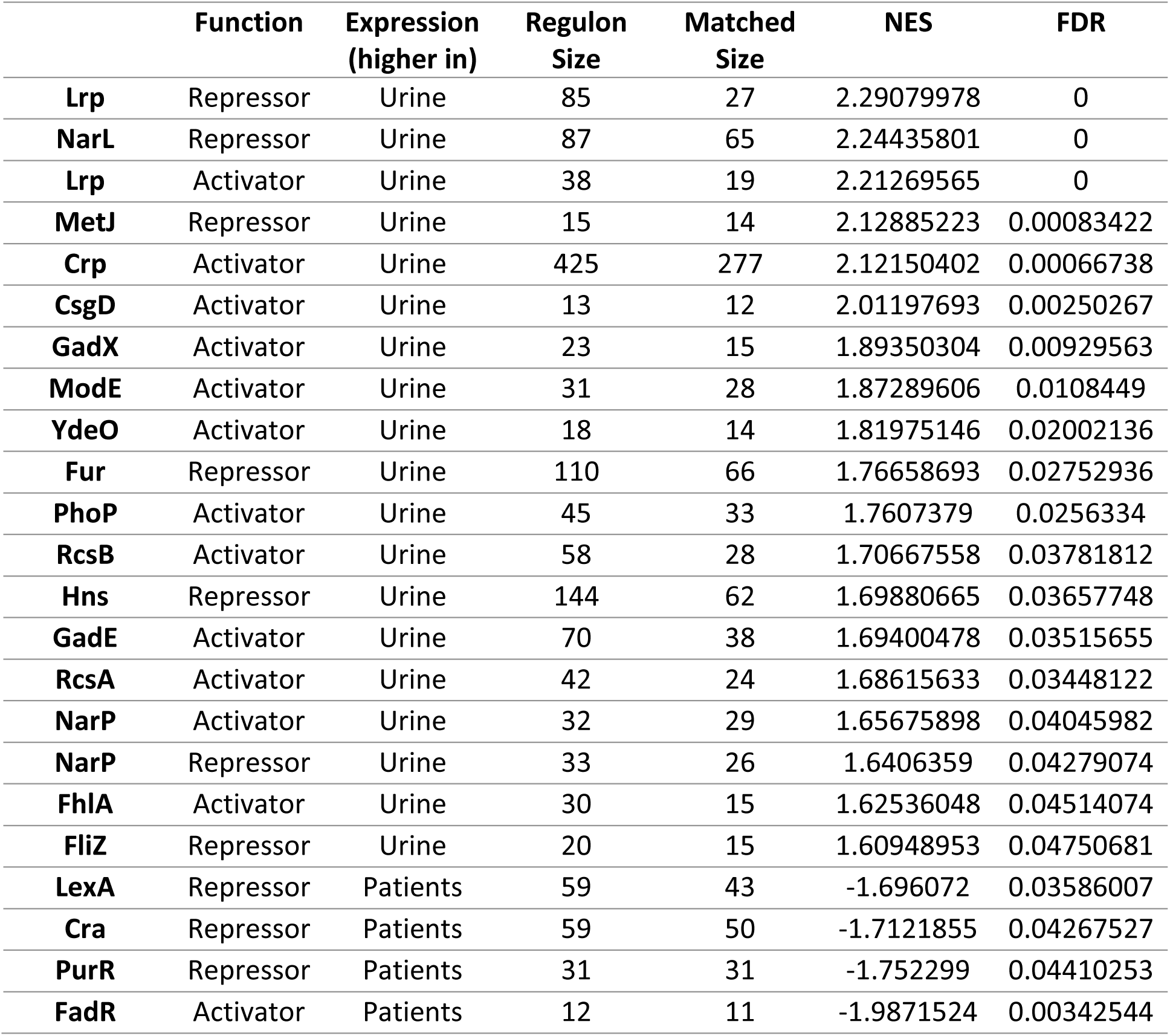
GSEA results^1^.

1 Gene sets found to be enriched in differentially expressed genes. For example, Lrp, Repressor indicates gene set repressed by Lrp (data obtained from RegulonDB 9.4). Expression indicates whether regulon expression was higher in patients of during *in vitro* culture in urine. Regulon size: number of genes in the gene set; Matched size: number of genes found in data set; NES: normalized enrichment score; FDR: false discovery rate.

